# Thymidylate synthase inhibitory drugs induce p53-dependent pathways differently

**DOI:** 10.1101/2025.09.02.673858

**Authors:** Eszter Holub, Anna Felföldi, Beáta G. Vértessy, Angéla Békési

## Abstract

Thymidylate synthase (TS) is a key enzyme in thymidylate biosynthesis and an established target of chemotherapeutics such as 5-fluoro-2’-deoxyuridine (5FdUR) and raltitrexed (RTX). Inhibition of TS disrupts the dUTP:dTTP balance, leading to uracil misincorporation, futile base excision repair cycles, DNA strand breaks, and ultimately cell death. Beyond its catalytic role, TS also binds RNA, autoregulating its own translation and interacting with transcripts such as p53 and c-myc, thereby linking TS activity to broader post-transcriptional regulatory networks. These interactions, together with regulation by miRNAs and lncRNAs, suggest that TS inhibition may provoke cellular responses extending beyond DNA metabolism. Non-coding RNAs, including miRNAs, snoRNAs, snRNAs, and lncRNAs, may play critical roles in shaping these outcomes. To dissect these mechanisms, we investigated the transcriptomic effects of TS inhibition in mismatch repair-deficient, UNG-inhibited HCT116 colon cancer cells treated with 5FdUR or RTX. Both drugs induced DNA damage responses and S-phase arrest, yet displayed distinct transcriptional signatures. Moreover, TS-RNA immunoprecipitation sequencing revealed direct RNA-binding targets of TS, highlighting its contribution as a post-transcriptional regulator. Our findings underscore the multifaceted impact of TS inhibition, linking enzymatic disruption to RNA-level regulation and revealing drug-specific differences in cellular responses.

## Introduction

Thymidylate synthase (TS) is one of the key enzymes of thymidine biosynthesis. It forms the sole *de novo* source of dTMP, a dTTP precursor, by transferring a methyl group from the 5,10-methylenetetrahydrofolate (MTHF) to the dUMP molecule. Cancer cells are actively dividing, thus targeting DNA synthesis is a common anticancer strategy, and inhibition of TS is widely used in chemotherapeutic treatment of solid tumors such as colon, breast, and head and neck cancers (1,2). TS is usually targeted by using substrate analogue molecules such as antifolates or fluoro-pyrimidines. Inhibition of the TS enzyme causes perturbation of the cellular dUTP:dTTP pool, which can lead to elevated uracil incorporation upon DNA synthesis, triggering base excision repair (BER) (3)). This repair pathway is initiated by the uracil-DNA-glycosylases (UDGs), among which the uracil-N-glycosylase (UNG) has the highest activity (4). Provided that the wrong dUTP:dTTP ratio persists, the repair cannot efficiently correct these uracils, which leads to hyperactive futile repair cycles, DNA strand breaks, and finally cell death (2). It has been shown that upon inhibition of UNG (with the expression of UGI protein), the genomic uracil level is dramatically elevated upon treatment with TS inhibitory drugs, 5-fluoro-2’-deoxyuridine (5FdUR) and raltitrexed (RTX), suggesting that these drugs efficiently perturb the cellular dUTP:dTTP ratio, which mostly leads to thymine replacing uracil incorporation (5,6). Among the several TS inhibitors that have been developed over the years, the nucleotide analogue 5FdUR and the antifolate RTX are the most specific ones (7,8). The active compound 5FdUMP is endogenously formed from 5FdUR, which creates a covalent ternary complex with TS and the MTHF molecule (9), while RTX is a competitive antifolate inhibitor (10).

It was shown that, besides having a crucial enzymatic role, TS is also able to bind to its own mRNA, autoregulating its protein level by translation inhibition (11,12). This phenomenon can provide a potential resistance mechanism against inhibitory drug treatments, as these drugs trap TS in the catalytically active conformation that is not compatible with RNA-binding (13). Hence, the TS protein tends to release its mRNA, allowing it to be translated; thus, the newly formed enzyme can counteract the drug effect (14). It was also shown that besides its own mRNA, TS can also bind to the mRNA of the tumor-suppressor, p53, and the oncogene, c-myc (15–17). Treatment with 5FU was found to cause increased TS mRNA level and also increased protein level of TS and p53 (18). The TS mRNA is additionally negatively regulated by different miRNAs (miR-197-3p, miR-203a-3p, miR-375-3p, miR-330-3p), which are targets of the MALAT1 lncRNA, usually up-regulated in cancers (19) and their transcription is also repressed by p53 (20). The TS mRNA is also the target of miR192/215-5p, which is induced by p53 (20). These studies suggest a complex regulatory network that might be influenced by the inhibitory drug treatments. As RNA-protein interactions are usually not predominantly sequence specific rather depend on the RNA secondary structure, these suggest a more general RNA regulatory role of TS with yet unidentified RNA binding partners.

Beyond the known mRNA targets of TS binding and regulation, non-coding RNAs can also be affected by TS-binding as well as by TS-inhibitory drug treatments. Short non-coding RNAs (<200–300 nucleotides) include small nuclear RNAs (snRNAs), small nucleolar RNAs (snoRNAs), and microRNAs (miRNAs). While snoRNAs primarily guide rRNA modification and maturation, and miRNAs mediate gene silencing through mRNA degradation or translational inhibition, the functions of other small RNAs, such as snRNAs, are more diverse. Long non-coding RNAs (lncRNAs), in contrast, are longer transcripts that do not encode proteins but regulate gene expression at multiple levels: transcriptionally, by modulating chromatin structure, and post-transcriptionally, often by sponging miRNAs to prevent degradation of their target mRNAs. They may also act as scaffolds in phase separation. Thus, TS inhibition is likely to trigger more complex cellular responses than previously recognized.

Previously, in mismatch repair (MMR) deficient, UNG-inhibited HCT116 colon adenocarcinoma cell-line, we published that both 5FdUR and RTX treatments result in S-phase arrested phenotype, and similar increase of γH2AX signal (6), which suggests that even though the main UDG is inhibited and mismatch repair is missing, these treatments eventually lead to the occurrence of DNA-strand breaks and the initiation of DNA damage response (DDR) signaling. Recently, we also detected differences between the two drug effects: 5FdUR, when applied in a high dose, was less effective than in a lower dose or any dose of RTX in UNG-inhibited, HCT116 cells, regardless of the MMR status.

To better understand the mechanism of action of these drugs and identify potential differences between them, we performed whole transcriptome sequencing from UNG-inhibited HCT116 cells upon these treatments. We could identify significantly up-regulated and down-regulated groups of RNAs, with some differences in the case of the two TS inhibitors. From these cells, we also performed TS-RNA-immunoprecipitation (TS-RIP) followed by sequencing, and addressed the contribution of the TS RNA-regulatory role to the whole cellular response.

## Materials and Methods

### Cell culturing

HCT116 UGI cells (6) were maintained in McCoy’s 5A medium (Thermo Fisher Scientific (Gibco), 16600082) supplemented with 10% FBS (Sigma, F9665-500ML) and 1% PenStrep (Thermo Fisher Scientific (Gibco), 15140122). Mycoplasma contamination was regularly checked. Cells were cultured in 6-well plates, treated with 100 nM or 20 µM of RTX or 5FdUR for 48 hours.

### RNA isolation

Drug-treated cells were harvested, and the total RNA pool was isolated using TRIzol reagent (ThermoFisher Scientific, 15596018) according to the manufacturer’s protocol. Briefly, cells were suspended in TRIzol reagent, incubated for 5 min at room temperature, and extracted with chloroform, then centrifuged at 12,000×g at 4 °C for 15 min. RNA-containing aqueous phase was collected, and RNA was precipitated with ice-cold isopropanol, washed with 75% ethanol, dried at 55 °C, and dissolved in RNase-free water. RNA sample quality was checked on a 1% agarose gel, and the concentration was measured using a NanoDrop 2000C (ThermoFisher).

### RNA-seq

Total RNA samples were used to establish stranded RNA sequencing libraries for short RNAs (below 200 nt), and long RNAs (above 200 nt), separately. Small and long RNA libraries were sequenced on a Novaseq6000 platform using Illumina 50 SE (20M reads), and 150 PE (150M reads), respectively. Library preparations, including rRNA depletion and QC, as well as the sequencing, were provided by Novogene Co., Ltd. (China) as a whole transcriptome sequencing service.

### RNA-seq analysis

From long RNA sequencing data, after quality check using fastQC (21), residual rRNA-related reads were removed in silico (script is provided in Supplementary Methods), then polyG, adapter, and low-quality sequences were trimmed using the fastp tool (22). Clean reads were aligned to the human reference genome GRCh38 (23) using HiSAT2 aligner (24). Transcriptome reconstruction was performed, and differential expression was analyzed using the Cufflinks package (25,26). Short RNA data were subjected to adapter and quality trimming using trimmomatic (27), then aligned to the GRCh38 reference genome using STAR (28). Read coverage was quantified on annotated mature miRNAs (hsa.gff3, mirBase, https://www.mirbase.org/download, (29,30)) and sRNAs, snoRNAs (GENCODE V34, https://www.gencodegenes.org/human/release_34.html, (31)) using the featureCounts tool (32), and compared using DESeq2 (33,34). Vulcano plots were created in R. Gene Set Enrichment Analysis (GSEA) was performed using the STRING database online analysis tool (35).

### qPCR validation of RNA-seq results

Reverse transcription-coupled quantitative PCR was performed to validate some of the findings of our RNA-seq analysis. Namely, we measured the gene expression levels of CDKN1A/p21, MDM2, and CCNG1/cyclin G, in addition to CNOT4 and PUM1, as reference genes. Specific primers for each target were used based on previous publications (36–39). The reverse transcription efficiency and the primer efficiency were checked for each target. 400 ng RNA was subjected to reverse transcription using the High-Capacity cDNA Reverse Transcription kit (Applied Biosystems, 4368814). Each 10 μl qPCR mix contained 2X MyTaq HS MIX (Bioline, BIO-25046), 20X EvaGreen Dye (Biotium, #31000), 500-500 nM reverse and forward primers, 0.15 μl reverse transcribed mix, and nuclease-free water. Each measurement was performed in technical triplicate and on three biological replicates with no-template control for each target in a CFX96 Touch Real-Time PCR Detection System (Biorad). The qPCR annealing temperature was 62 °C for 15 sec, which was followed by a 72 °C 30 sec elongation for 50 cycles. Melting curve analysis was also performed after each PCR reaction.

### TS-RNA-immunoprecititation (TS-RIP)

Cells were cultured on 15 cm Petri dishes, and treated with 100 nM RTX, 20 µM 5FdUR, or nothing (NT). After the 48-hour treatments and a wash with PBS, cells were harvested using a scraper and centrifuged (150×g, 21 °C), then resuspended in PBS for counting. After a second centrifugation, cells were resuspended in cytoplasmic extraction buffer (1 ml/10^7^ cells, composition: 20 mM TRIS, pH 7.4, 10 mM NaCl, 3 mM MgCl2, 0.5 mM dithiotreitol, 0.05% NP40, Roche Protease inhibitor tablet) and incubated on ice for 15 min with frequent mixing. Upon centrifugation (3000×g, 4 °C, 7 min), the supernatant was kept as the cytoplasmic fraction. The pellet was resuspended in nuclear extraction buffer (200 µl / 10^7^ cells, composition: 50 mM TRIS, pH 7.4, 150 mM NaCl, 50 mM NaF, 5 mM EDTA, 1 mM EGTA, 1 mM PMSF, 1% NP40, Roche Protease inhibitor tablet) and incubated on ice for 30 min with frequent vortexing. After centrifugation (16000×g 10 min 4 °C), the supernatant (nuclear extract) was collected, reunited with the cytoplasmic fraction, and supplemented with glycerol (final conc. 6.5%). Then these lysates were added to 55 μL pre-washed magnetic Protein AG beads (Thermo Scientific, Cat. no. 88803) and incubated at 4 °C for 2 h with constant rotation for pre-clearing. Beads were collected and used as a negative control; hence were resuspended in the same buffers and kept in the same conditions as the RIP samples. The pre-cleared lysate was subjected to TS-RNA-immunoprecipitation (RIP): first, overnight incubation with anti-TS-antibody (3-4.5 ug; Proteintech 15047-1-AP), then 2-hour incubation with washed magnetic protein A/G beads at 4 °C with constant rotation. Then, the beads were collected and washed three times with low-salt buffer (20 mM TRIS, pH 8.0, 150 mM NaCl, 2mM EDTA, 1% TritonX), and once with the washing buffer (DEPC-treated PBS, 0.05% Tween20). For elution, beads were incubated in 200 µl elution buffer (100 mM TRIS, 10 mM EDTA, 1% SDS, 1 µg/ml Proteinase K) at 60 °C for 30 min with frequent vortexing. 5% of the pre-cleared lysates were saved as input samples and incubated under the same conditions as the RIP samples, including the final Proteinase K treatment. The eluted solution was subjected to RNA isolation (Geneaid, VR050) following the manufacturer’s instructions. RNA concentration was measured with Qubit RNA BR assay kit (Thermo Fisher Scientific, Q10211) on a Qubit 4.0 (Thermo Fisher Scientific, Waltham, MA, USA). The RNA integrity in the samples was checked on 1% agarose gels compared to the input samples (Fig S1B). The presence of TS enzyme during the RIP was followed by Western blotting (Fig S1A). RNA samples (RIP and negative control) were sent for sequencing.

### TS-RIP-seq and data analysis

The libraries for Illumina sequencing were prepared using xGen RNA Lib Prep Kit (Integrated DNA Technologies, Iowa, USA). Briefly, 400 ng RNA was rRNA-depleted using RiboCop rRNA Depletion Kit for Human/Mouse/Rat V2 (Lexogen, Vienna, Austria). Thereafter, the RNA was fragmented and reverse-transcribed using random priming, then the cDNA was tailed and adapter-ligated. Finally, the libraries were amplified according to the manufacturer’s instructions. The quality of the libraries was checked on Agilent 4200 TapeSation System using D1000 Screen Tape (Agilent Technologies, Palo Alto, CA, USA), and the quantity was measured on Qubit 3.0 (Thermo Fisher Scientific, Waltham, MA, USA). Illumina sequencing was performed on the NovaSeq X Plus instrument (Illumina, San Diego, CA, USA) with 2 × 151 run configuration. The whole procedure of library preparation and sequencing was performed as a service of the Hungarian Centre for Genomics and Bioinformatics, Szentágothai Research Centre at the University of Pécs, Hungary.

Sequencing data were processed by the same pipeline as the longRNA sequencing data described earlier. TS-RIP samples were compared to the corresponding RIPctr samples. In addition, TS-RIP enrichment was also calculated for exons. For this, all human exons annotated in GenCode V34 were filtered for those that have a detectable average read coverage in the transcriptome sequencing. Read coverage and log_2_(fold change) tracks were calculated using the deeptools package (40).

### Western blot

Samples were subjected to SDS-PAGE using a 12% polyacrylamide gel, followed by transfer to PVDF membrane (Millipore; IPVH09120) in 10 mM CAPS 15% EtOH pH 12, 350 mA 3 h. The membrane was blocked using 5% milk powder in PBS-T. Primary antibodies were added in 5% BSA TBS-T (anti-TS-ab. 1:3000;(Proteintech 15047-1-AP); anti-p53-ab. (DO-1) 1:2000 (Santa Cruz sc-126); anti-p21-ab 1:1000 (Cell signaling Technology #2947); anti-actin-ab. 1:2500, (Sigma A1978); anti-γH2AX-ab. 1:750 (Millipore 05-636) and incubated at 4°C overnight (ON). After several washing steps, HRP-conjugated secondary antibodies were added (1:10,000) and incubated for 2 h at RT. After several washing steps with PBS, visualisation of the membrane was performed using Immobilon Western Chemiluminescent HRP Substrate (Millipore) on a ChemiDoc MP Imaging System (Bio-Rad). Western blot images were analyzed and further processed with ImageLab software (Bio-Rad).

## Results

### RTX and 5FdUR treatments influence the cellular gene expression profiles differently

To better understand and describe the differences between these two TS inhibitory drugs (RTX and 5FdUR) effects on the cellular response, we performed whole transcriptome sequencing (Fig 1., data are available at GEO). Relative expression levels and differential expression were determined as described in the Methods and Supplementary Methods. We found a comparable amount of down-regulated and up-regulated genes in the case of protein-coding RNAs (Fig 1A) and short RNAs (Fig 1C), in contrast with long-non-coding RNAs (lncRNAs, Fig 1B), where a prominent bias was detected towards down-regulation.

**Fig 1.**
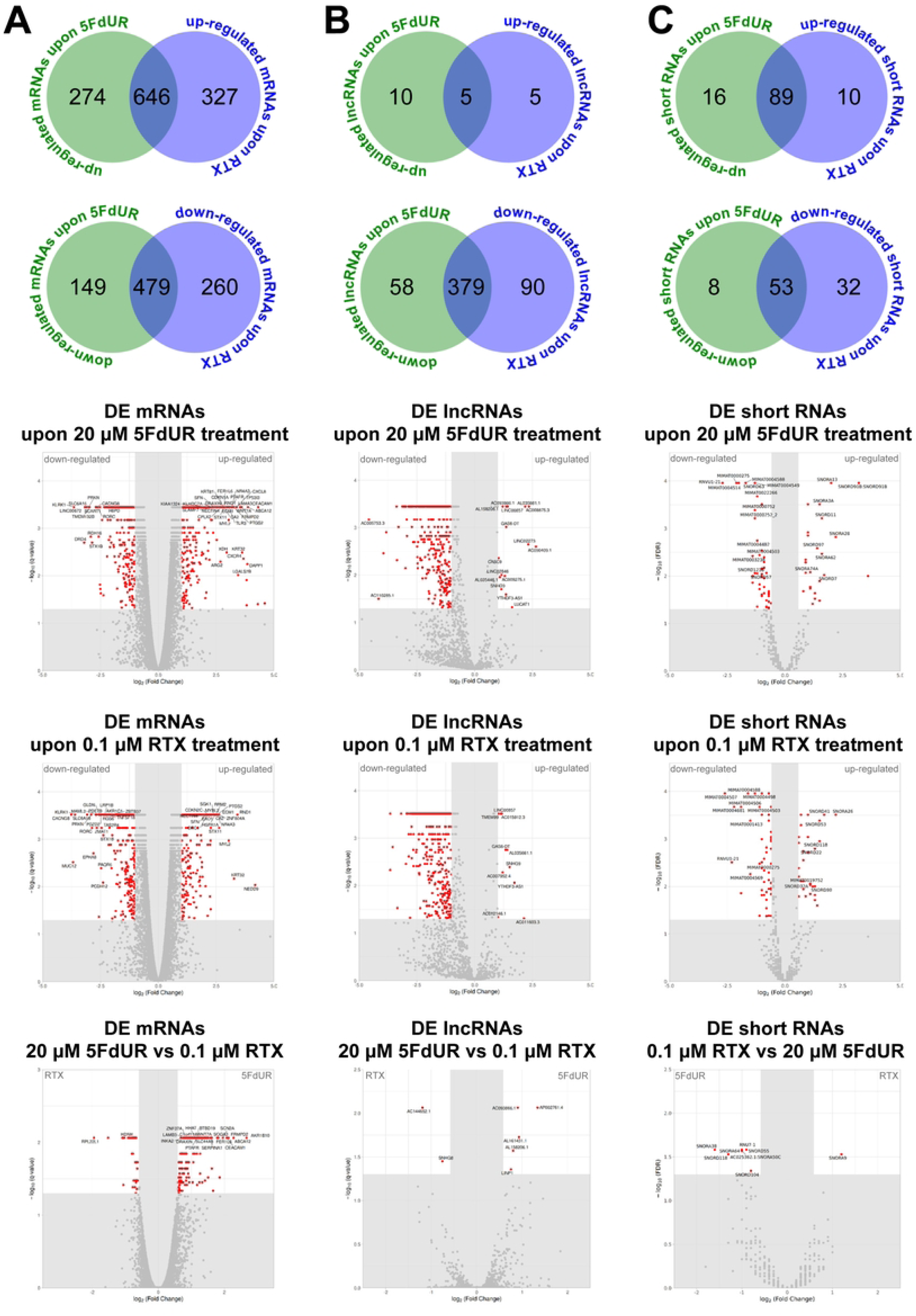
Differentially expressed (DE) genes upon RTX and 5FdUR treatments. Differential expression was calculated based on whole transcriptome sequencing data using the Cufflinks package (25,26) with two biological replicates. For significance, the following filters were applied to the long RNAs: minimum fold change +/-2 (or +/-1.5 in the case of the comparisons of RTX and 5FdUR samples), q-value < 0.05, and mean FPKM > 1000. While filters for the short RNAs were: minimum fold change +/-1.5 FDR < 0.05; mean counts > 50. **Significantly up-or down-regulated mRNAs (A), lncRNAs (B), and short RNA (C).** Venn diagrams (on the top) show the number of significantly up-regulated or down-regulated RNAs upon 0.1 μM RTX (blue) or 20 μM 5FdUR (green) treatments. Vulcano plots show the distribution of DE RNAs (significant red or non-significant gray dots) either for the two treatments compared to the non-treated (NT) sample, or for comparison of the two drugs (as indicated on the plots). The most outstanding DE mRNAs based on q-value and log_2_FC, as well as all upregulated lncRNAs, are labeled by their gene symbols. For short RNAs, the gene names are used as labels.

In general, more differentially expressed (DE) genes were identified upon RTX, as compared to 5FdUR-treatment (cf. Venn diagrams in Fig 1). However, upregulation of non-coding RNAs (lncRNAs, sRNAs, snoRNAs, and miRNAs) more frequently occurs upon 5FdUR as compared to RTX. In addition, the fold change values of these increased expressions are also higher in general in the case of the 5FdUR, as compared to the RTX-treatment, and this tendency is true for the protein-coding RNAs as well (cf. Volcano plots in Fig 1). To evaluate the significance of these drug-specific differentially regulated expressions, we compare directly the two drug-treated samples (cf. bottom Volcano plots in Fig 1). These plots also confirm a stronger upregulation caused by 5FdUR, which potentially indicates additional pathways activated by 5FdUR-treatment as compared to the RTX.

### Differentially expressed non-coding RNAs in the case of RTX and 5FdUR

In our data, up-regulated short RNAs were mostly snoRNAs, while among the down-regulated ones, several miRNAs were also found (volcano plots in Fig 1C). When we compared the two drugs, we found that only SNORA9 showed significant RTX-bias, while SNORA38, SNORD118, SNORA64, RNU7-1, SNORA50C, SNORD55, and SNORD104 showed significant 5FdUR-bias. Interestingly, no miRNA was found to be significantly differently expressed when comparing the drug treatments. However, besides the several down-regulated snoRNAs, 6 and 33 miRNAs were found significantly downregulated in the case of 5FdUR-and RTX-treatment, respectively. We subjected the top three of these miRNAs that show the highest expression to a network using the online platform of miRNet (41) (https://www.mirnet.ca/miRNet/upload/MirUploadView.xhtml) (Fig S2). KEGG enrichment on RNAs found in the miRNA network revealed that the p53-related network is targeted by these selected, high-level, but down-regulated miRNAs in both cases. Additionally, in the case of RTX treatment, cell cycle regulation and stress response were also enriched terms.

As most of the lncRNAs were down-regulated upon treatment with RTX or 5FdUR (Volcano plots in Fig 1B), we focused on the few up-regulated ones. LINP1, AL158206.1, AL161431.1, AC093866.1, and AP002761.4 show a significant 5FdUR-specific increase compared to RTX, while the gene expression level of AC144652.1 and SNHG8 are significantly elevated upon RTX treatment.

### Drug-induced expression changes confirm S-phase-arrested and DNA damage-induced phenotype, and impaired transcriptional activity

To gain a better understanding of the processes and pathways involved in the cellular response upon treatment with 5FdUR or RTX, a GSEA was performed on the significantly up-regulated and down-regulated groups of protein-coding genes using the online platform of the STRING database (35) (https://string-db.org, Fig 2 and Table 1). We found that in the case of the commonly up-regulated genes, terms of cell-cycle regulation, cellular response to DNA damage, DNA repair, DNA replication, and extracellular region were the most enriched terms. This is in good agreement with the strong S-phase arrest detected upon both treatments and elevated gamma-H2AX signals (6). Consistently, members of the p53 transcriptional network, ATR and ATM signaling pathways, cellular senescence, and apoptosis signaling were also enriched upon both drug treatments.

**Fig 2.**
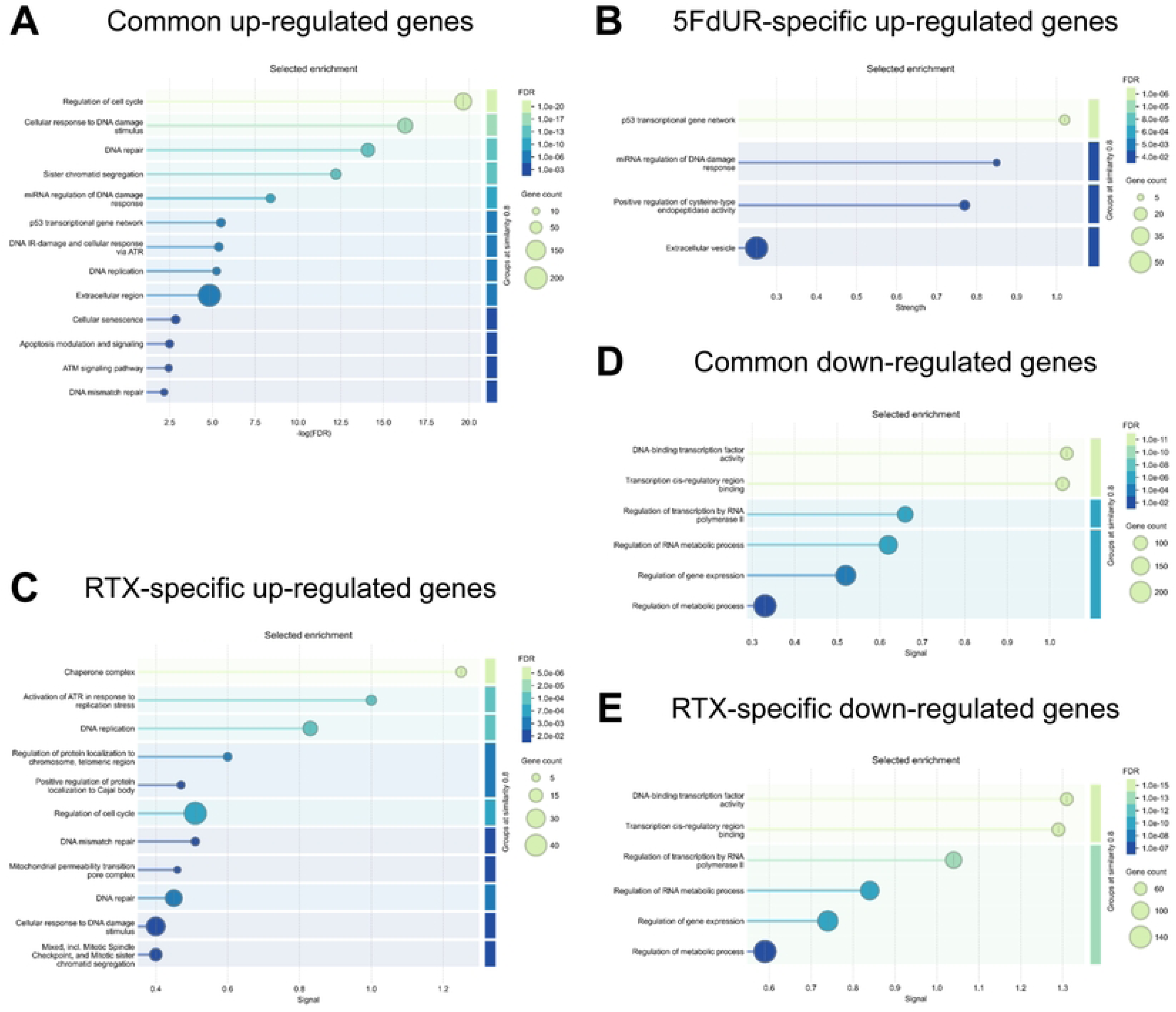
Gene Set Enrichment Analysis (GSEA) of differentially expressed RNAs. GSEA was performed using the online analysis tool of the STRING database (35). Selected enriched terms are shown on the enrichment maps measured in gene sets of up-or down-regulated genes, either by both treatments or just specifically by 0.1 μM RTX, or 20 μM 5FdUR treatments, as indicated on the panels. No enriched terms were found in the case of 5FdUR-specific down-regulated genes. The significantly up-and down-regulated genes are listed in the source data file. Permanent links for the corresponding STRING networks and network statistics are given in Table 1.

**Table 1:**
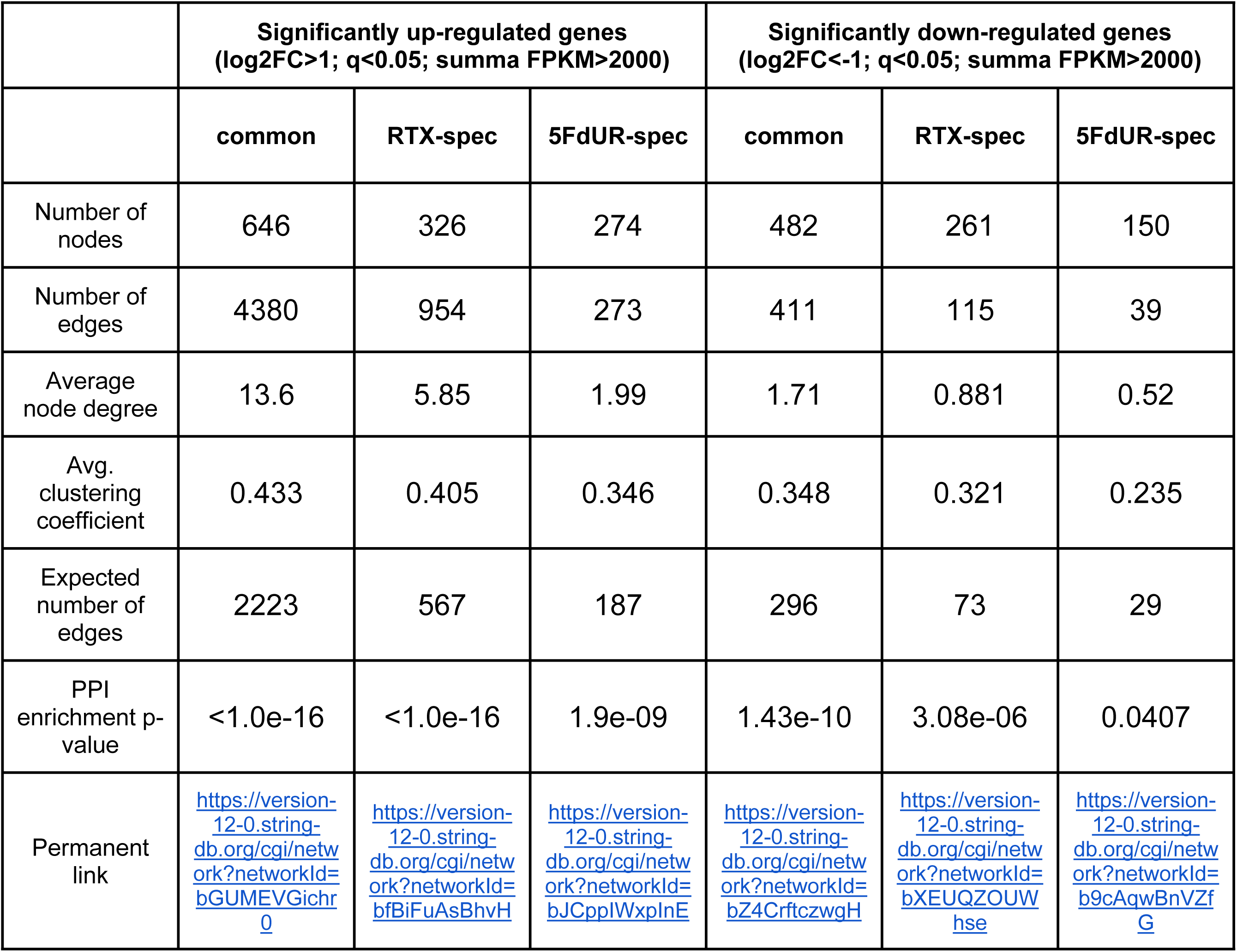
Network statistics and permanent links from GSEA (STRING). Interactions in the STRING database were filtered for “medium confidence level”, and only “textmining”, “experiments”, and “databases” were considered as valid sources.

Above the common upregulated processes, 5FdUR-specific up-regulated genes showed further enrichment in the components of the p53 transcriptional and signaling network, as well as in transcripts of proteins related to extracellular processes and miRNA regulation of DNA damage (Fig 2B). While RTX-specific induction mainly affects processes related to replication stress, protein folding, cell-cycle regulation, and coupled chromatin organization and assembly, as well as mitochondrial permeability (Fig 2C).

Genes down-regulated upon both drug treatments were enriched in processes involved in the regulation of gene expression were particularly abundant, such as transcription factor activity, regulation of transcription by RNA-Pol-II, and regulation of RNA metabolic processes (Table 1 and Fig 2DE). These terms were also among the most enriched in the case of RTX-specific down-regulation; however, no further enrichment was found among the genes specifically down-regulated upon 5FdUR treatment.

These observed biases in the case of up-regulated genes in the case of 5FdUR treatment towards the p53 network and extracellular elements, and in the case of RTX treatment towards the cell cycle and replication-related and repair processes, complemented with RTX-specific bias towards the down-regulation of components of gene expression, report differences in the cell response upon treatment with these TS inhibitors.

### Altered activation of p53-regulatory network upon RTX and 5FdUR treatments

To better visualize how the p53 regulatory networks are affected by drug-induced expression changes, first, we highlighted these genes and TP53 itself on the Volcano plots, also labeling those genes that show significant differences in the RTX and 5FdUR comparison (Fig 3A). Interestingly, TP53 is not induced at the mRNA level; however, many of its target genes are significantly induced. The p53-mediated regulation is usually initiated within the DDR, and deciding on cell death or survival, has multiple possible outputs. To understand the drug-specific differences better, we mapped the differential expression level (RTX compared to 5FdUR) to the KEGG network of p53 signaling pathway (hsa04115) (Fig 3B). It is immediately visualized that apoptosis-related subpathways are different for the two drugs: induction of reactive oxygen species (ROS) might be more involved in 5FdUR-mediated cytotoxicity, while SIVA1, the only one with RTX-biased expression, inhibits the anti-apoptotic BCL2 proteins. 5FdUR-biased induction of angiogenesis and metastasis inhibitory factors (including serine protease inhibitors PAI (SERPINE1) and Maspin (SERPINB5), and other extracellular proteins, thrombospondin 1 (TSP1) and CD82 antigen (KAI)) are in good agreement with the enriched cellular component of “extracellular region” (cf. Fig 2B). The p21 (CDKN1A, a key cell-cycle regulatory protein) has an extremely high expression level that is further induced by both drugs, but with a significant bias towards 5FdUR. These differences would suggest increased cytotoxicity of 5FdUR treatment, but that is not the case. However, we have to point to the 5FdUR-biased induction of MDM2, providing a negative feedback loop for p53 at the protein level.

**Fig 3.**
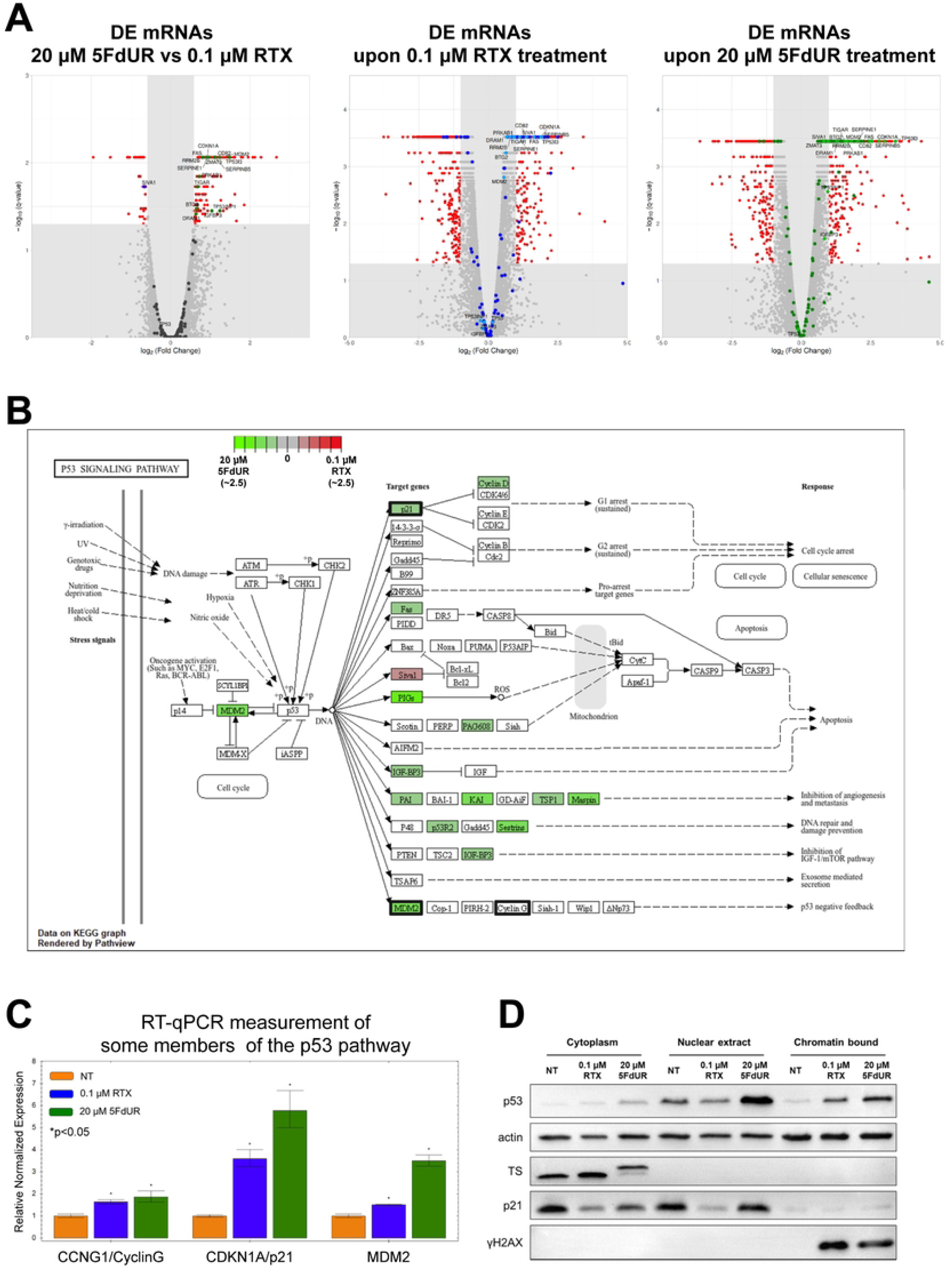
The p53 pathways are more induced upon 5FdUR treatment. A: On the volcano plots (same as in Fig 1A), components of the “p53 signaling pathway” (KEGG, hsa04115) and the “p53 transcriptional gene network” (WikiPathways, WP4963) are marked with dark gray in the non-significant zone, and colored when they show significant differences (RTX blue, 5FdUR green). Those that are significantly different in the comparison of RTX and 5FdUR (left plot) are labeled with their gene symbols on all three plots. On the middle and right volcano plots, where drug-induced DE RNAs are colored, those p53 network components that are also significantly different between the two drugs are represented with lighter shades of the same color. **B:** Drug-specific differential expression levels are projected to the “p53 signaling pathway” (KEGG, hsa04115). Colorbar from green to red on the top follows the log_2_(fold change) up to 2.5 in both directions. **C:** Induction of CCNG1 (cyclin G), CDKN1A(p21), and MDM2 at the mRNA level, measured with RT-qPCR in the non-treated (NT), 20 μM 5FdUR, or 0.1 μM RTX 48 h treatment (n=3). **D:** Expression and subcellular localization of p53, TS, p21, and γH2AX proteins in the case of TS inhibitory drug treatments.

The above analysis of RNA-seq data vividly demonstrates that the p53 network was induced much more upon 5FdUR treatment than upon treatment with RTX (cf. Fig 3AB). To validate these high-throughput results, we measured the gene expression level of 3 genes in the p53 network with RT-qPCR (Fig 3C). We chose the CDK inhibitor p21 encoded by the CDKN1A gene, and two negative regulators of p53, the E3 ubiquitin ligase MDM2 (MDM2 gene), and the cyclin G encoded by the CCNG1 gene. We found that the mRNA level of p21/CDKN1A was highly elevated upon 5FdUR treatment and to a lesser extent upon RTX treatment (fold change values are 5.78 ± 0.78; 3.60 ± 0.36, respectively). Gene expression level of MDM2 was also found to be strongly increased upon 5FdUR treatment and modestly increased upon RTX treatment (3.50 ± 0.25; 1.50 ± 0.03). In the case of cyclin G similarly slight change was found upon both drug treatments (fold change values are: 1.87 ± 0.23; 1.64 ± 0.09, respectively). These results are in good agreement with our RNA-seq analysis, further confirming our findings.

As many p53 target genes were elevated but p53 mRNA expression level was found to be unchanged (cf. Fig 3A), we checked p53 protein level upon the drug treatments by Western blotting (Fig 3D). Indeed, in the nuclear extract fraction of 5FdUR-treated cells, a massive p53 signal increase was detected. In the case of RTX-treated cells, p53 showed only a weak induction as compared to the NT sample and only within the chromatin-bound fraction (Fig 3D). These results suggest additional regulatory mechanisms, either at the translational level or at the protein stabilization level of p53, that are stronger in the 5FdUR sample than in the RTX-treated one. This occurs despite the 5FdUR-specific induction of MDM2 transcript (cf. Fig 3ABC). We also checked the DNA damage marker, γH2AX, which also showed a similarly immense signal increase in the chromatin-bound fraction for both drug treatments, indicating comparable intensity of DDR. TS protein could only be detected in the cytoplasmic fractions, and in accordance with the RNA-seq results, it is slightly induced upon RTX treatment, but not in the 5FdUR-treated sample. Instead, a band duplication can be observed here that is related to the strong ternary complex of TS, 5FdUMP, and the folate co-substrate (9). In addition, we also checked p21, the protein product of the other validated drug-induced gene, CDKN1A. Interestingly, while the mRNA level of p21 was highly elevated upon the drug treatments, the protein level was the same or even decreased in the cases of 5FdUR and RTX, respectively (Fig 3D). This surprising result indicates a strong translation inhibitory mechanism that is stronger upon RTX treatment than 5FdUR.

### The effects of drug treatments on TS RNA-binding ability

As TS was described as a moonlighting protein, also possessing RNA-binding and translation-inhibiting ability (11,12), we assumed that drug treatments could affect these functions, which might explain the observed discrepancies between the mRNA and protein levels of p53 and p21. Hence, we performed RNA co-immunoprecipitations using anti-TS antibody (TS-RIP, scheme in Fig 4A), and sequenced the TS-RIP and the corresponding control IP (RIPctr, pull-down without specific antibody) samples. We could pull down both the p53 and the TS mRNAs, but surprisingly, they were not markedly enriched when their relative abundances were compared between the TS-RIP and RIPctr samples (Table 2). Interestingly, drug treatments increased these enrichments in most cases except for the TS mRNA in the 5FdUR sample.

**Fig 4.**
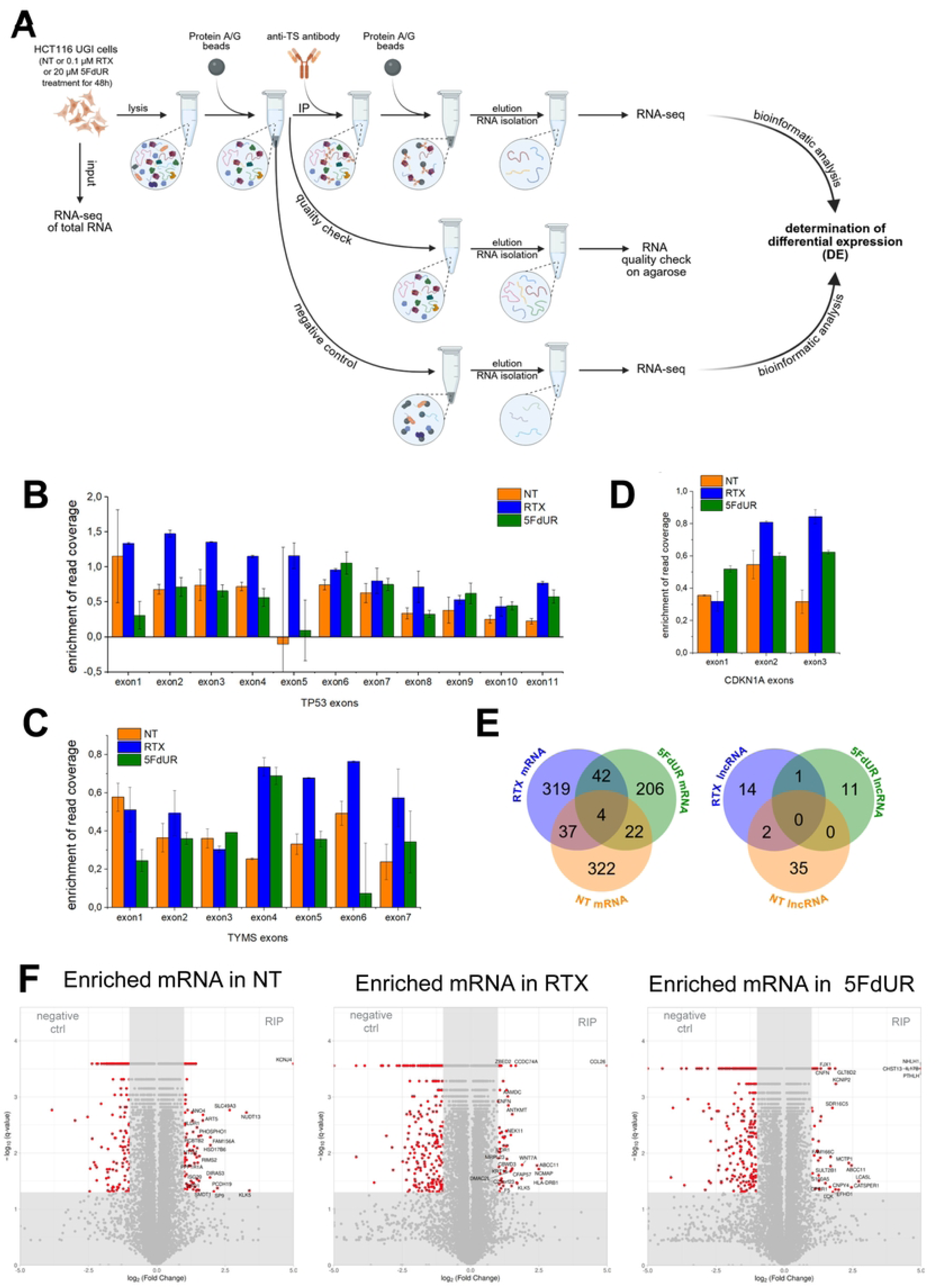
Several RNAs are bound to thymidylate synthase. A: Workflow of TS-bound RNA immunoprecipitation (TS-RIP), which was followed by next-generation sequencing. Average read coverage values were calculated separately for exons, and TS-RIP values were compared to the corresponding negative control (RIPctr) in non-treated (NT), 0.1 μM RTX-treated, or 20 μM 5FdUR-treated samples. These TS-RIP enrichment profiles along the exons of TP53/p53 (**B**), TYMS/TS (**C**), and CDKN1A/p21 (**D**) were visualized on bar graphs. Standard deviation is calculated based on pseudoreplicate data. **E:** Venn diagrams of TS-bound mRNA (left) or lncRNA (right) in case of 0.1 μM RTX, 20 μM 5FdUR, or not treated (NT) samples. Enrichment was considered to be significant when log_2_FC>0.585, q-value<0.05, and the mean FPKM > 500. **F:** Volcano plots of TS-RIP compared to the corresponding RIPctr samples in case of non-treated (NT), 0.1 μM RTX, or 20 μM 5FdUR treatments. Labelling of the most enriched genes was done based on log_2_FC and q-value.

**Table 2:** TS-RIP enrichment data for the selected genes in the three samples: NT, RTX, and 5FdUR, calculated with the cuffdiff tool as described in Supplementary Methods.

| sample | RNA | FPKM<br>RIPctr | FPKM in TS-RIP | $\log_2$ (fold<br>change) | q-value |
| --- | --- | --- | --- | --- | --- |
| NT | TP53 | 94520 | 94081 | -0.007 | 0.932 |
| **RTX** | **TP53** | **88631** | **115828** | **0.386** | **0.00028** |
| **5FdUR** | **TP53** | **98913** | **115836** | **0.228** | **0.00031** |
| NT | TYMS | 126651 | 125622 | -0.012 | 0.855 |
| **RTX** | **TYMS** | **241156** | **277632** | **0.203** | **0.00028** |
| 5FdUR | TYMS | 200383 | 195808 | -0.033 | 0.548 |
| **NT** | **CDKN1A** | **159164** | **181984** | **0.193** | **0.00026** |
| **RTX** | **CDKN1A** | **362117** | **478637** | **0.402** | **0.00028** |
| **5FdUR** | **CDKN1A** | **1.04966e+06** | **1.27427e+06** | **0.280** | **0.00031** |

Furthermore, changes in the enrichment profiles could also be observed when average log_2_(fold change) values are calculated for distinct exons (Fig 4BCD). In the case of TS mRNA, the highest enrichment was detected on the first and sixth exons in the non-treated (NT) sample (Fig 4C). While a significantly increased enrichment was observed on the 4th exon upon both drug treatments, on the 1st and 6th exons, a 5FdUR-specific decrease was detected compared to NT. This, in accordance with the overall enrichment values (cf. Table 2), might indicate that the TS mRNA is less bound to the TS enzyme upon 5FdUR than RTX treatment. In the RTX sample, enrichment was induced on the 4th, 5th, 6th, and 7th exons, while remaining similar on the first, 2nd, and 3rd exons of TS mRNA as compared to the NT sample (Fig 4C). This suggests that upon RTX treatment, the TS mRNA might bind to the TS protein stronger than in the case of NT or 5FdUR samples. As the RTX treatment caused an elevated TS protein level (cf. Fig 3D), the translation-inhibitory effect of this interaction is most probably fine-tuned by the actual mode of interaction that is influenced by the exposure to RTX. These results suggest that interaction at the beginning of the TS mRNA might play a role in its translational inhibition, as TS-RIP enrichment was decreased in this region upon drug treatment, which is in agreement with the protein induction. In contrast, interaction at the end of the mRNA might contribute to RNA stability.

In the case of the p53 mRNA, another RNA known to interact with the TS protein, we could identify a similar trend as in the case of TS mRNA (Fig 4B). Treatment with 0.1 μM RTX caused an increase in enrichment of read coverage on almost every exon of *TP53* except exons 1, 5, and 7, when it was similar to NT. On the other hand, treatment with 20 μM 5FdUR only caused increased enrichment on the 6th, 9th, and 11th exons, and even decreased enrichment of read coverage was detected on the 1st exon compared to NT. These findings suggest that p53 mRNA is bound to the TS enzyme more weakly in the case of 5FdUR treatment and stronger upon RTX treatment, presumably allowing it to be translated into protein in the former case. This aligns with our findings that p53 mRNA level is unchanged in our total RNA-seq data upon 5FdUR treatment, while its protein level is significantly elevated. This also proposes that these two TS inhibitory drugs might influence TS-mRNA binding capacity differently.

In the case of p21 mRNA encoded by the gene CDKN1A, a weak enrichment was observed in the non-treated sample that was increased upon drug treatments, especially upon RTX (cf. Table 2). These tendencies were true for exons 2 and 3, while exon 1 showed increased enrichment only upon 5fdUR treatment (Fig 4D).

### Thymidylate synthase binds to several RNAs *in vivo*

We could identify several mRNAs and a few lncRNAs bound to TS, which were not reported before (Fig 4E and F). From this data, we could also detect that treatments with 5FdUR or RTX do not similarly influence TS RNA binding capacity. While only 42 mRNAs were found common upon both treatments but not in the non-treated sample, 319 and 206 mRNAs were RTX-or 5FdUR-specific, respectively, while 322 were NT-specific. In the non-treated and RTX-treated samples, 37 mRNAs were found in both; 22 mRNAs were found common in the non-treated and 5FdUR-treated samples, while 4 mRNAs were found in all samples. We subjected the lists of enriched protein-coding genes upon RTX, 5FdUR, or no treatment to GSEA (Table 3) and found significant enrichment of membrane organelles in all cases (Table 3). In the NT sample intrinsic component of membrane, the integral component of membrane, the bounding membrane component, and the organelle membrane were the most enriched gene ontology (GO) terms. Upon RTX treatment, mitochondrial membrane, striated muscle thin filament, and cytoplasm GO terms were additionally present, while upon 5FdUR treatment, the endoplasmic reticulum membrane GO term was found to be enriched as well. No functional enrichment was detected among the groups of mRNAs common between NT and RTX or NT and 5FdUR, and only the Human T-cell leukemia virus 1 infection KEGG pathway was enriched in the mRNAs common between the drug treatments.

**Table 3:**
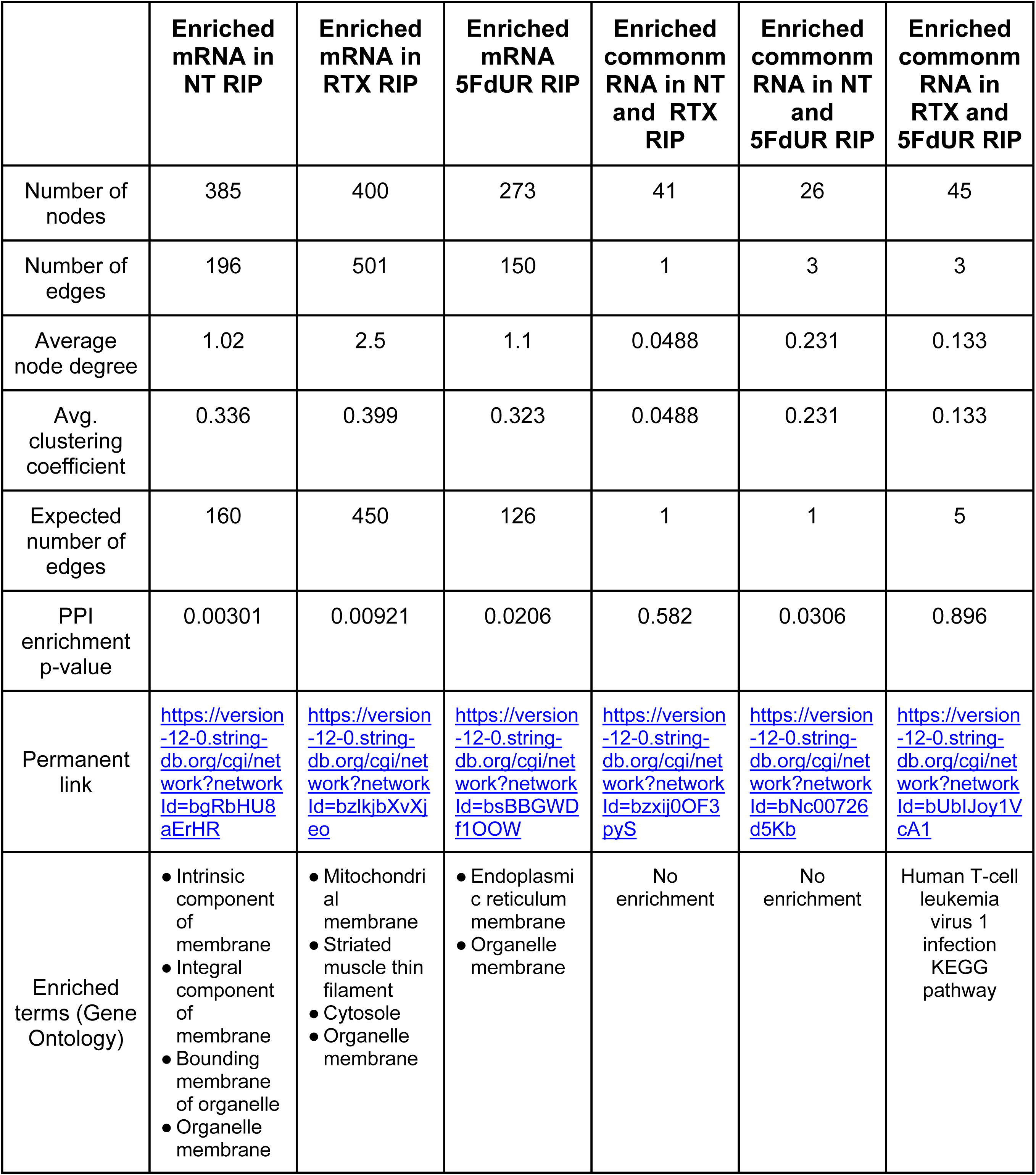
GSEA results and STRING network statistics of enriched mRNAs bound to TS without treatment or upon treatment with 0.1 μM RTX or 20 μM 5FdUR.

In general, a magnitude less lncRNA was identified as TS-bound than mRNA (Fig 4E). Additionally, less than half the amount of lncRNA was found in the drug-treated cases, which might be due to the overall downregulation of lncRNAs upon treatment with TS inhibitors. Among the TS-bound lncRNAs, AC080112.2 was found to be common between RTX and 5FdUR, 0 between NT and 5FdUR, and 2 (AC027031.2, AL109627.1) between NT and RTX. The AC027031.2 lncRNA is proposed to take part in ferroptosis (42), while about AL109627.1, there is no publication available yet. The lncRNA AC080112.2 is not yet characterised, and its possible role in the cellular context is unidentified. The binding of TS to lncRNAs logically cannot influence their translation as they do not code for proteins; rather, it might modulate the stability of these lncRNAs.

In summary, in the case of RTX treatment, a similar amount of mRNA was found, but less lncRNA compared to NT, while generally less TS-bound RNA (mRNA and lncRNA) was identified upon 20 μM 5FdUR treatment. Our findings ultimately suggest that the covalent ternary complex formed upon the binding of the 5FdUR metabolite 5FdUMP disrupts the interaction between the TS protein and RNA more than the binding of the antifolate RTX, which in some cases might even enhance the TS protein-RNA interaction.

## Discussion

Thymidylate synthase inhibitory drugs are widely used in chemotherapeutic treatments; however their exact mechanism of action is not yet fully understood. Here, we provide deeper insight into the cellular response at the RNA level initiated upon treatment with antifolate RTX or base-analogue 5FdUR. We also present evidence that although these drugs target the same enzyme, there are differences in the induced cellular response between them. We also hypothesize that some of these differences might originate from TS’s role in mRNA regulation. Accordingly, we show that TS protein interaction with its own mRNA and p53 is altered differently in the case of the two inhibitors. Additionally, we also identified previously undescribed RNA-interacting partners of TS protein that differ upon inhibition with 5FdUR or RTX.

Functional enrichment analysis of mRNA significantly up-regulated upon TS inhibitory drug treatment revealed that the cell cycle and DNA damage response, as well as the p53 signaling network, were affected upon both drug treatments, particularly upon treatment with 5FdUR. RNA metabolism and transcription-related processes were enriched among the drug-induced significantly down-regulated protein-coding genes, which were even enriched in the RTX-specific group. Interestingly, no functional enrichment was found in the case of 5FdUR-specific down-regulation.

We also detected that most lncRNAs are downregulated in the drug-treated cells compared to the non-treated control. This down-regulation of lncRNAs might be the consequence of the general down-regulation of transcription activity. Additionally, we hypothesize that this down-regulation of gene expression upon TS inhibitory drug treatments might be due to the changes in the dynamics of the transcriptome. We think that the generation of new transcripts is less frequent upon drug treatments compared to NT, while the ones detected as up-regulated RNAs might be stabilized somehow, maybe as a result of cell response to the stress caused by these drugs, thus the degradation of these RNAs is postponed. In this regard, the few up-regulated lncRNAs might have some significance in the cellular response as well.

RTX-specific up-regulated lncRNAs, AC144652.1 and SHNG8, were reported as potential prognostic biomarkers for cancer, with high expression correlating with high risk and worse overall survival. While lncRNA AC144652.1 and its interacting network are relatively uncharacterized, its potential as a prognostic marker via its possible involvement in cuproptosis-related processes in the case of head and neck squamous cell carcinoma was recently proposed (43). On the contrary, SHNG8 and its regulatory network are much better described. Its high expression was observed in various cancers and was associated with tumour growth and worse prognosis. Upon silencing lncRNA SNHG8, its role to sponge miR-384, miR-634, miR-588, miR-152, miR-491, and miR-149-5p was compromised, thus cell proliferation, invasion, autophagy, and increased resistance were attenuated (44). Ultimately, the upregulation of lncRNAs AC144652.1 and SHNG8 might be beneficial for cancer cell survival upon chemotherapy.

In the case of 5FdUR-specific up-regulated lncRNAs, the image is a bit more complex. High expression level of AL161431.1 and AP002761.4 was associated with worse prognosis and poor survival, as they promote cell proliferation (45,46). On the other hand, the high expression of LINP1 increased sensitivity to radiotherapy by enhancing the non-homologous end joining pathway, resulting in erroneous repair, which can disrupt genes and their regulation, finally leading to cell death (47). Low expression level of lncRNA AP002761.4 was associated with worse survival in a bioinformatic study researching disulfidptosis-related lncRNAs in the case of acute myeloid leukaemia (48), suggesting that high expression in our case might be unfavorable for tumor cell survival. LncRNA AL158206.1 was found to be up-regulated 4 hours post high-dose ionising irradiation of primary skin fibroblasts, regardless of the donor’s cancer history (49). In this study, weighted-coexpression analysis revealed that in the case of AL158206.1, the most adjacent mRNAs were CDKN1A, MDM2, TIGAR, BTG2, and HSPA4L; accordingly, signal transduction by p53 class mediator was the most enriched when functional enrichment analysis was performed on differentially expressed mRNAs. This suggests that a similar cellular response is initiated after treatment with TS-inhibitory drugs, particularly with 5FdUR, as these mRNAs are also found to be upregulated in our study. It is also supported by the high γH2AX signal in our cells following treatment with RTX or 5FdUR, which is usually a sign for DNA double-strand breaks, typically caused by ionising radiation.

Functional enrichment analysis of down-regulated miRNA networks revealed the p53-signaling pathway and cell cycle, which were found to be enriched terms in the case of up-regulated mRNAs upon TS inhibitory treatment, revealing that the miRNA-based negative regulation of p53 pathway processes in our cells is not present.

As the p53 pathway was found to be significantly up-regulated upon both drug treatments compared to the non-treated sample, we also compared the enrichment upon RTX and 5FdUR treatments. 5FdUR biased up-regulation was found in the p53 pathway, which was further confirmed with RT-qPCR measurement of the mRNA levels of p53 regulated genes p21, MDM2 and cyclin G. We also checked the protein level of p53 as its gene expression was not significantly changed upon treatments, so we hypothesised that the differential expression level of p53-regulated genes must be due to elevated protein levels of p53. We found a similar protein level of p53 in the chromatin-bound fraction upon both drug treatments, which was higher than in the NT sample. Additionally, a prominent increase in p53 protein level was also detected in the nuclear extract only in the case of 5FdUR treatment. Contrary to the changes detected at the mRNA levels of p21, the protein level was found to be unchanged or even less upon TS inhibitory drug treatments compared to the NT. This contravention might be explained by the post-transcriptional regulation of the p21 mRNA via the calreticulin protein that inhibits its translation to protein (50). We checked the gene expression level of calreticulin in our RNA-seq data and found an increase upon both drug treatments, which was stronger in the case of RTX, which matches our observation on the p21 protein level. As lncRNA SPUD, generated from the CDKN1A gene via alternative polyadenylation, is also known to regulate p21 expression by competing with its mRNA in binding to calreticulin (51). We found that SPUD expression level was overall low and did not change significantly upon treatments, thus the detected p21 protein expression is probably the result of calreticulin-mediated translational inhibition. Our results show that the p53 pathway is initiated upon both TS inhibitory treatments. In addition, we could also detect differences between the induced RNAs upon treatments with RTX or 5FdUR, which might contribute to the biased activation of the p53 pathway and, after all, to altered cell survival upon 20 μM 5FdUR treatment (manuscript under revision).

A potential modulating factor might be TS and its ability to bind RNA (just like calreticulin), which might contribute to different cellular responses upon TS inhibitory drug treatments. To investigate whether there are any differences upon treatment with the drugs in TS RNA binding capacity, we performed RNA immunoprecipitation via the TS protein that was followed by sequencing. We could detect differences in the RNA binding in TS when we compared the enrichments of read coverages of different exons of its own mRNA and the mRNA of p53, two already identified binding partners. Our results revealed that upon treatment with 5FdUR, the mRNA of p53 is less bound to the TS protein than in the case of RTX treatment or NT. This result provides an explanation on how the p53 protein level is strongly elevated without changes in its mRNA level only upon 5FdUR treatment. We could also identify that its own mRNA, the mRNA of p53 and c-myc (the known RNA-interacting partners of TS) were not among our strongest hits. We could identify several new RNAs that were bound to TS, which, interestingly, were all enriched in elements related to membranes when GSEA was performed. We could also detect a few lncRNAs among the TS-associated RNAs whose stability might be influenced upon TS binding. At the same time, the interaction with TS might interfere with the post-transcriptional regulatory role of lncRNAs, which possibility needs further investigation to rule out.

In conclusion, our results report differences in cellular response at the RNA level between TS-inhibitory drugs, RTX, and 5FdUR. We also present that the RNA regulatory function of TS is much wider than previously reported, and it is differently influenced upon inhibition with RTX or 5FdUR. Our findings provide additional insight into these drugs’ mechanisms of action, enabling us to better understand the cellular responses arising upon treatment with RTX or 5FdUR, which might even contribute to resistance upon treatment with these inhibitory drugs.

## Acknowledgements

We acknowledged Gábor Tusnády for providing access to computational capacity. We acknowledge the ENCODE Consortium (<u>ENCODE Project Consortium, 2012</u>) and the ENCODE production laboratory(s) generating the particular dataset(s), as well as the mirBASE, the STRING, and the mirNET databases and free online tools supporting our research. We also say thanks for the customization of their NGS service to Péter Urbán and Bence Gálik at the Hungarian Centre for Genomics and Bioinformatics, Szentágothai Research Centre at the University of Pécs, Hungary.

---

**Supplementary Fig S1:**
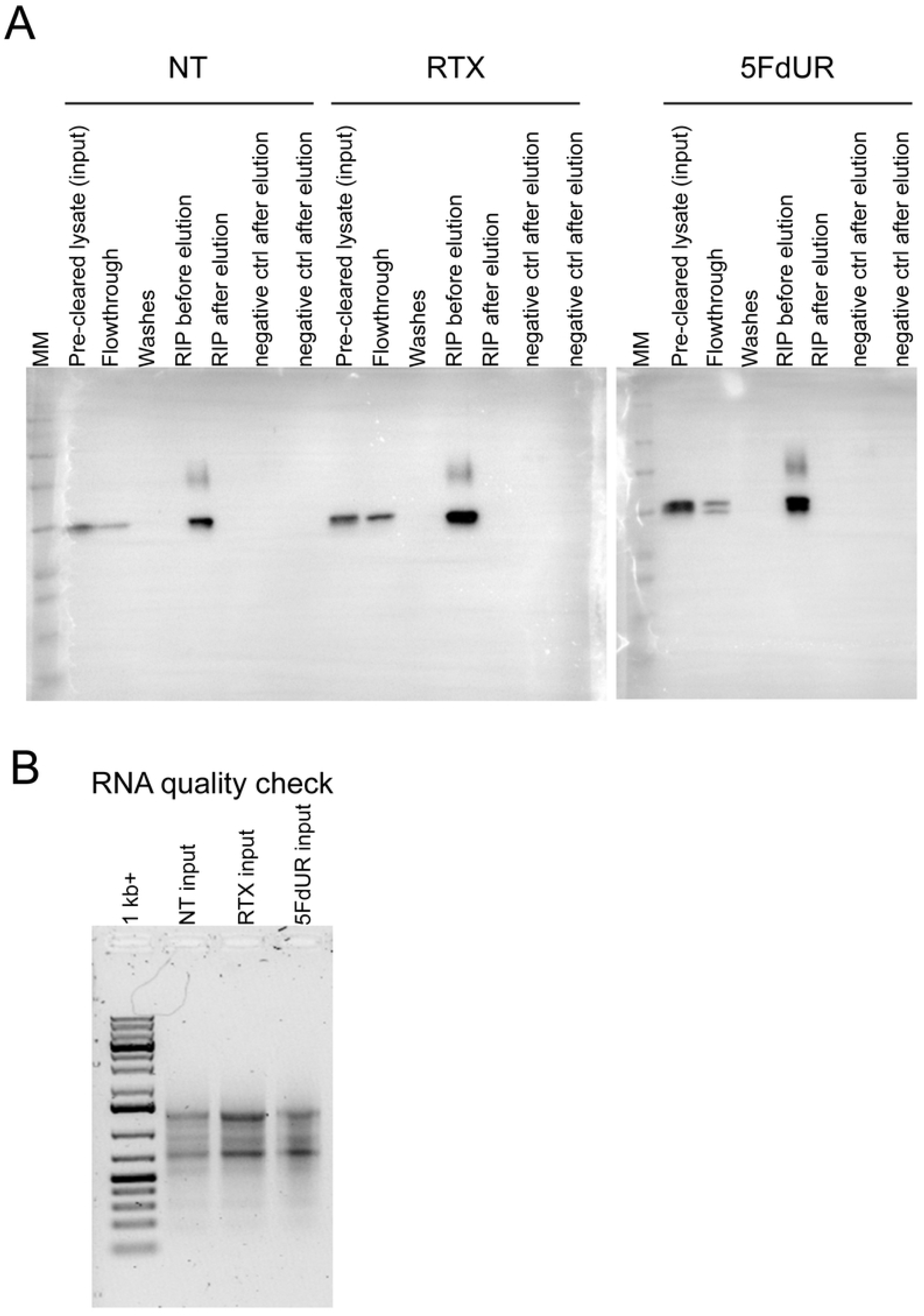
TS-RIP workflow followed by Western blotting and RNA sample quality control. A Western blotting on TS-RIP fractions. MM: GRS Protein Marker MultiColour (GRISP Research Solutions, Porto, Portugal), “RIP before elution” indicates the antibody-conjugated beads that successfully recruited TS protein from the pre-cleaned lysates (NT, RTX, 5FdUR samples as indicated at the top. “Negative ctrl” samples represent protein A/G agarose incubated with lysates without anti-TS antibody. B: RNA samples isolated from the saved portion of the input samples and incubated under the same conditions as the TS-RIP samples were. The two main bands correspond to the abundant rRNAs, which were preserved during the long immunoprecipitation procedure.

**Supplementary Figure S2:**
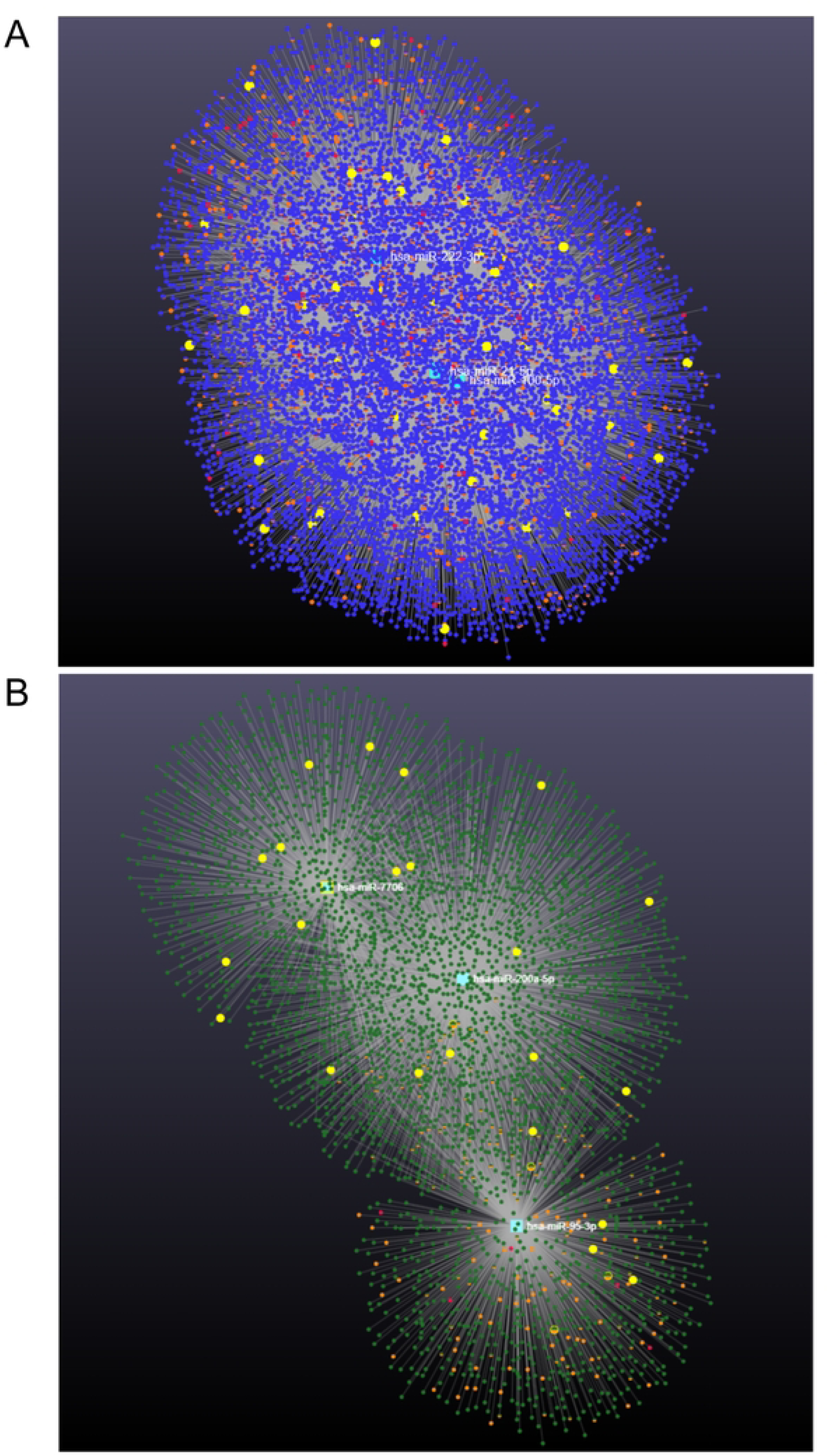
ceRNA network of the top 3 down-regulated miRNAs. The top three miRNAs in RTX samples were hsa-miR-21-5p (mean counts ∼8M, log2(FC) =-0.5, FDR=0.0078), hsa-miR-100-5p (mean counts ∼ 1.5M, log2(FC) =-0.6, FDR=0.0016), and hsa-miR-222-3p (mean counts ∼ 100K, log2(FC) =-0.4, FDR=0.0347); and in 5FdUR samples were hsa-miR-200a-5p (mean counts∼44K, log2(FC) =-0.7, FDR=0.048), hsa-miR-7706 (mean counts∼4.8K, log2(FC) =-0.8, FDR=0.0325), and hsa-miR-95-3p (mean count∼4.5K, log2(FC) =-0.7, FDR=0.023). The target search was done against TarBase9.0, lncRNA (red dots), and circRNA (orange dots). mRNA targets are blue (RTX) or green (5FdUR), while query miRNAs are enlarged squares in cyan. A GSEA was performed on the same online platform against KEGG pathways, and p53-related miRNA targets are enlarged in the network and marked with yellow.

